# The shape of aroma: measuring and modeling citrus oil gland distribution

**DOI:** 10.1101/2022.04.14.488418

**Authors:** Erik J. Amézquita, Michelle Y. Quigley, Tim Ophelders, Danelle Seymour, Elizabeth Munch, Daniel H. Chitwood

**Affiliations:** Department of Computational Mathematics, Science & Engineering, Michigan State University, East Lansing, MI, USA; Department of Horticulture, Michigan State University, East Lansing, MI, USA; Department of Mathematics, Michigan State University, East Lansing, MI, USA; Department of Mathematics and Computer Science, TU Eindhoven, Eindhoven, The Netherlands; Department of Information and Computing Sciences, Utrecht University, Utrecht, The Netherlands; Department of Botany & Plant Sciences, University of California, Riverside, CA, USA

**Author notes:** To whom correspondence should be addressed: Dr. Daniel H. Chitwood, 1066 Bogue St, East Lansing, MI 48824, Dr. Elizabeth Munch, 428 S Shaw Ln, East Lansing, MI 48824.

**Keywords:** directional statistics, citrus, oil glands, mathematical biology, data science, shape

## Abstract

Citrus come in diverse sizes and shapes, and play a key role in world culture and economy. Citrus oil glands in particular contain essential oils which include plant secondary metabolites associated with flavor and aroma. Capturing and analyzing nuanced information behind the citrus fruit shape and its oil gland distribution provides a morphology-driven path to further our insight into phenotype-genotype interactions.
We investigated the shape of citrus fruit of 51 accessions based on 3D X-ray CT scan reconstructions. Accessions include all three ancestral citrus species, accessions from related genera, and several interspecific hybrids. We digitally separate and compare the size of fruit endocarp, mesocarp, exocarp, and oil gland tissue. Based on the centers of the oil glands, overall fruit shape is approximated with an ellipsoid. Possible oil gland distributions on this ellipsoid surface are explored using directional statistics.
There is a strong allometry along fruit tissues; that is, we observe a strong linear relationship between the volume of any pair of major tissues. This suggests that the relative growth of fruit tissues with respect to each other follows a power law. We also observe that on average, glands distance themselves from their nearest neighbor following a square root relationship, which suggests normal diffusion dynamics at play.
The observed allometry and square root models point to the existence of biophysical developmental constraints that govern novel relationships between fruit dimensions from both evolutionary and breeding perspectives. Understanding these biophysical interactions prompt an exciting research path on fruit development and breeding.

**Societal Impact Statement:** Citrus are intrinsically connected to human health and culture, including preventing human diseases like scurvy, and inspiring sacred rituals. Citrus fruits come in a stunning number of different sizes and shapes, ranging from small clementines to oversized pummelos, and fruits display a vast diversity of flavors and aromas. These qualities are key in both traditional and modern medicine and the production of cleaning and perfume products. By quantifying and modeling overall fruit shape and oil gland distribution, we can gain further insight into citrus development and the impacts of domestication and improvement on multiple characteristics of the fruit.

## 1 Introduction

*“Fairest of all God’s trees, the orange came and settled here*,

… *Alone and unmoving you stand: how can one not admire you? Deep-rooted, hard to shift: truly you have no peer!*

*Alert to this world’s ways you hold your ground, unyielding against the vulgar tide*.*”*

—from Qu Yuan (340–278 BC?), “In Praise of the Orange-Tree (Ju song)” (Hawkes, 1985).

Citrus fruits and leaves have played a fundamental role across multiple aspects of human history including the development of modern nutrition and medical sciences. The aromatic and medicinal properties of mandarins and oranges have inspired delicate poetry since ancient times (Tseng, 1999; Vovin, 2016). Etrog citrons represent “the fruit of a goodly tree” during the Sukkot celebrations in the Jewish community (Isaac, 1959). The bael tree is considered sacred and it is generally grown near Hindu temples (Sharma et al., 2007). The fruits, peels, and leaves of diverse citrus have been used as traditional medicine for millennia for a diverse array of maladies (Mahomoodally and Mooroteea, 2021; Shrestha and Dangol, 2019). Sour oranges and lemons inspired the first modern clinical trials in the 18th and 19th centuries to determine the causes and cure of scurvy thus paving the way to the eventual isolation and synthesis of the first vitamin, vitamin C. (Baron, 2009; Magiorkinis et al., 2011).

Currently there is a rising trend in global citrus production, with more than 143 million tonnes produced in 2019 alone (FAO, 2021). Citrus production is valued for more than 3.3 billion US dollars in the US alone. (NASS, 2021). Citrus derived products are vital for other multi-billion dollar industries as well, from orange juice in the food industry, to essential oils in the perfume and cosmetics industry (Spreen et al., 2020). Essential oils in particular are extracted for their aromatic, flavoring, medicinal, and preservation properties useful in a variety of contexts (Mahato et al., 2019).

Before any human intervention, current paleobotanical evidence suggests that the common ancestor of citrus species originated more than 8 million years ago in the triangle defined by modern day northeastern India, northern Myanmar, and northwestern Yunnan (Talon et al., 2020). As monsoons weakened in southeastern Asia and climate transitioned to drier conditions, citrus radiated and diversified over the next 5 million years across the southeast Asian peninsula, Australia, New Caledonia, the western Indian coast, and even Japan (Wu et al., 2018). Early civilizations in India and China domesticated some of these species and their interspecific hybrids, even as early as during the Xia Dynasty (2100-1600 BC) in Southern China (Deng et al., 2020). Through tribute, trade, and invasion, different cultures contributed to spread many of these citrus across the rest of the world over the next 3000 years (Langgut, 2017).

Citrus species are sexually compatible and their ability to hybridize, combined with constant displacement and cultivation in multiple environments, produced a diversity of admixed accessions with a vast array of phenotypic traits (Gmitter et al., 2020; Luro et al., 2017; Wu et al., 2021). Asexual propagation is common in citrus and the interaction between grafted individuals has led to novel phenotypes, including through the formation of graft chimeras, conglomerations of cells that originated from separate zygotes (Caruso et al., 2020). The first reported plant chimera, known as Bizarria, arose from a fortuitous graft junction of a Florentine citron and a sour orange in 1674 (Nati, 1674). Since then, chimeras have proved to be more common than originally thought, transforming our perception of the genetic heterogeneity of individuals and its impact on plant development and phenotype (Frank and Chitwood, 2016).

A phenotype of particular interest is shape. Specific combinations of shape features are used to distinguish diverse citrus varieties, and have motivated various citrus taxonomic systems (Ollitrault et al., 2020). Leaf shape has been used to distinguish pummelos from sweet oranges among other different citrus genotypes and their respective environment interactions (Iwata et al., 2002). Root architecture is indicative of soil deficiencies for sour orange rootstocks (Mei et al., 2011). Morphological traits, such as fruit size and oil gland density, are used to infer genetic similarities between various mandarin cultivars (Pal et al., 2013). Oil gland size, structure and distribution are associated with the fruit development of navel oranges (Knight et al., 2001) and grapefruits (Voo et al., 2012).

Here we study the shape of citrus fruits and fruits from close citrus relatives based on 3D X-ray CT (computed tomography) scan reconstruction of 166 different samples comprising 51 different accessions, including samples of the three fundamental citrus species (*C. medica, C. reticulata*, and *C. maxima*), accessions from related genera (*P. trifoliata* and *F. margarita*), and several interspecific hybrids. First, using the power of X-rays and image processing, we compared volume ratios between different tissues, including exocarp, endocarp, and oil gland tissue. Second, since citrus oil glands contain essential oils which include plant secondary metabolites associated with flavor and aroma, we examined the number of individual oil glands, their density, and their overall distribution across all fruits. We determine that the average distance between neighboring oil glands follows a square root model, which indicates that gland distribution might follow normal diffusion dynamics (Vlahos et al., 2008). Finally, based off a point cloud defined by the center of all individual oil glands, we model the fruit shape as an ellipsoidal surface, a sphere with its three main axes shrunk or stretched accordingly. Once the glands are considered points on an ellipsoid, we are able to apply multiple tools from directional statistics (Ley and Verdebout, 2017; Mardia and Jupp, 1999; Pewsey and García-Portugués, 2021), which allows us to study and infer possible statistical distributions on spherical surfaces. As an example of this mathematical machinery, we test whether the oil glands either follow a uniform or symmetric distribution across the fruit surface. To the best of our knowledge, the shape of citrus has not been explored with similar scanning technologies, nor has it been analyzed with ellipsoidal and directional approximations. This morphological modeling will allow us to set a new exciting path to explore further the phenotype-genotype relationship in citrus.

## 2 Materials and Methods

### 2.1 Plant material and scanning

We selected 51 different accessions of citrus and citrus relatives with diverse fruit morphologies and geographical origins for our analysis. Fruits were sampled from a single tree for each selected accession maintained in the University of California Riverside Givaudan Citrus Variety Collection. 166 different individuals in total were sent for scanning at Michigan State University in December 2018 (Figure 1(a); Table S1.) These 166 samples were arranged into 63 raw scans, one scan per citrus variety containing all the replicates. Pummelos and citrons samples were scanned individually due to the fruit size. The scans were produced using the North Star Imaging X3000 system and the included efX software, with 720 projections per scan, at 3 frames per second and with 3 frames averaged per projection. The data was obtained in continuous mode. The X-ray source was set to a current of 70 µA, voltage ranging from 70 to 90 kV, and focal spot sizes ranging from 4.9 to 6.3 µm. The 3D reconstruction of the fruit was computed with the efX-CT software, obtaining final voxel sizes ranging from 18.6 to 110.1 µm for different scans (Figure 1(b); Table S2.)

**FIGURE 1.**
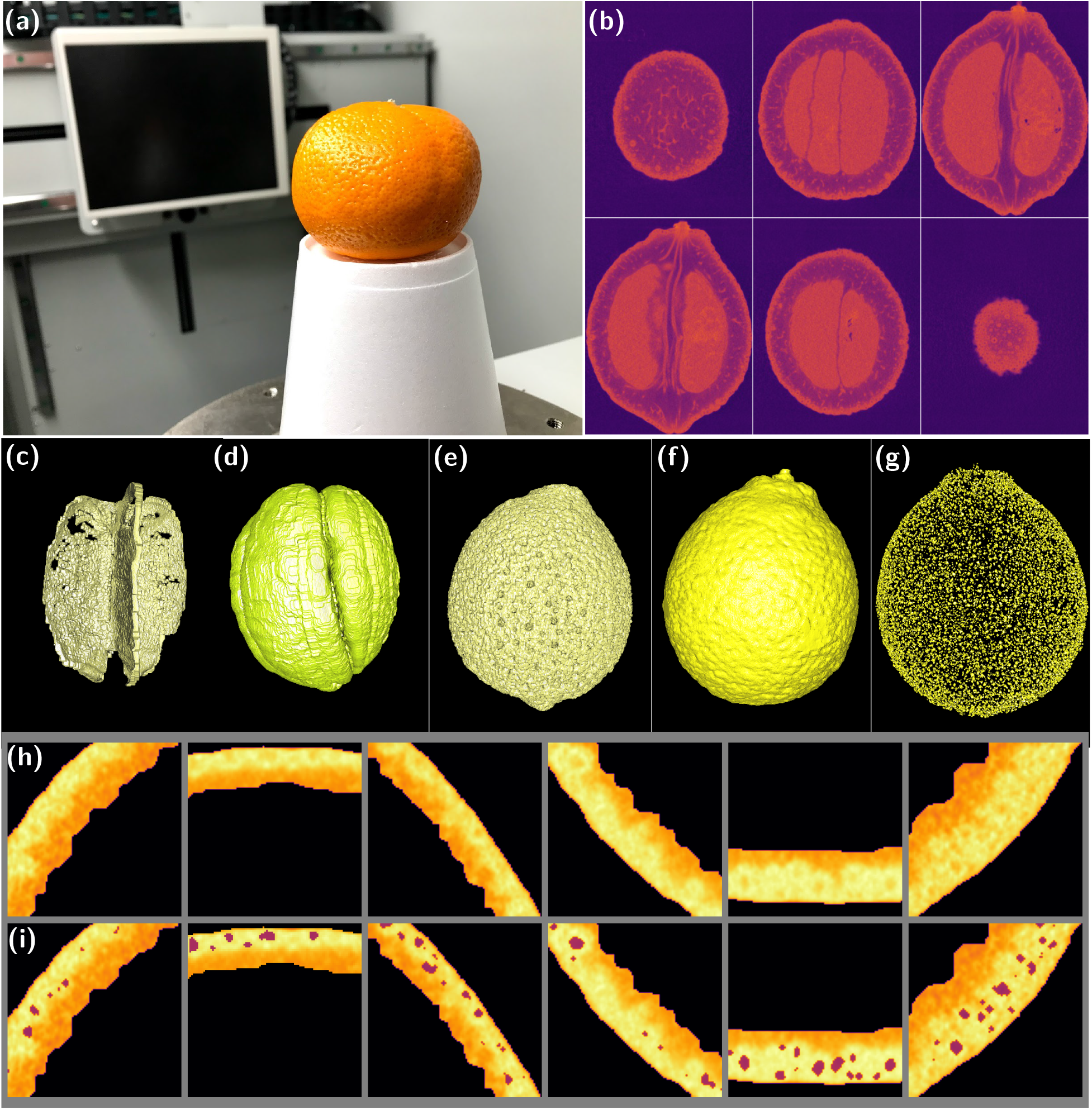
Citrus scanning and image processing. (a) A diverse collection was scanned using X-ray CT technology. (b) Slices of a raw scan. The image processing steps involved segmenting individual fruits and removing air and other debris. Then, individual tissues for each fruit were segmented such as the (c) central spine, (d) endocarp, (e) mesocarp, (f) exocarp, and (g) oil glands. (h) Close-up of some X-ray slices of the exocarp. (i) Same figure as above, with the segmented oil gland tissue darkened for emphasis. A Willowleaf sour orange is used as an example for figures (b)–(i).

(The air and debris were thresholded out of each raw scan, and individual replicates segmented into separate images. Based on density and location, for each fruit we further segmented 3D voxel-based reconstructions of its central column, endocarp, mesocarp, exocarp, and oil glands (Figure 1(c)–(g)). The center of each oil gland was calculated as the center of mass of the voxels composing such gland. An in-house scipy.ndimage-based python script was used to process the images for all fruits and their tissues. These were later visually inspected to verify their correctness. All the Chinese box oranges (*Severinia buxifolia*) scans were discarded due to their poor quality. To highlight nuanced differences among certain citrus groups, scans were partly split into sensible clusters of morphological interest (Table 1).

**TABLE 1.** Selected citrus groups and varieties. **N** equals the number of pseudo-replicates. The full list of citrus fruits and relatives scanned is found in Table S1. Names according to the University of California Givaudan Citrus Variety Collection (CVC). Asterisk denotes not available.

| CVC Name | Scientific Name | N | - | CVC Name | Scientific Name | N |
| --- | --- | --- | --- | --- | --- | --- |
| **Kumquats** |  |  |  |  |  |  |
| Nagami kumquat | *F. margarita* (Lour.) | 3 |  |  |  |  |
| **Lemons and lemon hybrids** |  |  |  |  |  |  |
| Limon Real | *C. excelsa* Wester | 4 |  | Lamas lemon | *C. limon* L.Burm.f. | 3 |
| Interdonato lemon | *C. limon* L. Burm.f. | 3 |  | Volckamer lemon | *C. volkameriana* | 3 |
| Frost nucellar Eureka lemon | *C. limon* L.Burm.f. | 3 |  |  |  |  |
| **Mandarins and mandarin hybrids** |  |  |  |  |  |  |
| Scarlet Emperor mandarin | *C. reticulata* Blanco | 3 |  | * | *C. reticulata* | 3 |
| Lee mandarin | *C. reticulata* Blanco | 3 |  | Som Keowan | *C. reticulata* Blanco | 3 |
| Koster mandarin | *C. reticulata* Blanco | 3 |  | Cleopatra mandarin | *C. reshni* hort.ex Tanaka | 3 |
| Beledy mandarin | *C. reticulata* Blanco | 4 |  | Fremont mandarin | *C. reticulata* Blanco | 3 |
| USDA 88-2 (Lee X Nova) | *C. reticulata* Blanco | 3 |  | Kinkoji Unshiu | *C. neo-aurantium* | 3 |
| **Microcitrus** |  |  |  |  |  |  |
| Australian finger lime | *M. australasica* | 4 |  |  |  |  |
| **Papedas** |  |  |  |  |  |  |
| * | *C. hanayu* Siebold | 3 |  | Kalpi, Nogapog | *C. webberii* | 2 |
| Makrut lime | *C. hystrix* | 4 |  |  |  |  |
| **Pummelos and pummelo hybrids** |  |  |  |  |  |  |
| Tahitian X Star Ruby | *C. paradisi* Macfadyen | 3 |  | Pomelit hybrid | *C. maxima* (Burm.) Merr | 3 |
| Kao Pan pummelo | *C. maxima* (Burm.) Merr. | 3 |  | Hassaku (Beni) hybrid | *C. hassaku* hort.ex Tanaka | 3 |
| Egami Buntan pummelo | *C. maxima* (Burm.) Merr. | 3 |  |  |  |  |
| **Sour oranges and sour orange hybrids** |  |  |  |  |  |  |
| Willowleaf sour orange | *C. aurantium* var. *salicifolia* | 3 |  | Standard sour orange | *C. aurantium* L. | 3 |
| Konejime hybrid | *C. neo-aurantium* | 4 |  | Olivelands Sour orange | *C. aurantium* L. | 3 |
| **Sweet oranges** |  |  |  |  |  |  |
| Cutter Valencia nucellar | *C. sinensis* L. Osbeck | 3 |  | Shamouti orange | *C. sinensis* L. Osbeck | 3 |
| Parent Washington navel | *C. sinensis* L. Osbeck | 3 |  | Argentina sweet orange | *C. sinensis* L. Osbeck | 3 |
| Cara Cara pink navel | *C. sinensis* L. Osbeck | 3 |  |  |  |  |
| **Trifoliates and trifoliolate hybrids** |  |  |  |  |  |  |
| Little-leaf trifoliolate | *P. trifoliata* (L.) Raf. | 3 |  | Rubidoux trifoliolate | *P. trifoliata* (L.) Raf. | 4 |
| C-35 citrange | *X Citroncirus* spp. | 3 |  | Carrizo citrange | *X Citroncirus* spp. | 5 |
| Swingle citrumelo | *X Citroncirus* spp. | 5 |  |  |  |  |

### 2.2 Allometric relationships

The total volume of fruits and their separate tissues was measured from the scans, as well as the number of individual oil glands. We studied the allometric relationships between these measurements; that is, the relative size of different tissues with respect to each other. These relationships in plants often follow a power law, so all the measurements were first log-transformed (Niklas, 2004) and a reduced major axis linear regression (Smith, 2009) was fitted considering all fruits. The slope, intercept, and *R*^2^ correlation coefficient were recorded (Figures 2, S1). The distribution of the residues was compared against a normal distribution to determine the adequacy of the linear fit (Figures 3, S2).

**FIGURE 2.**
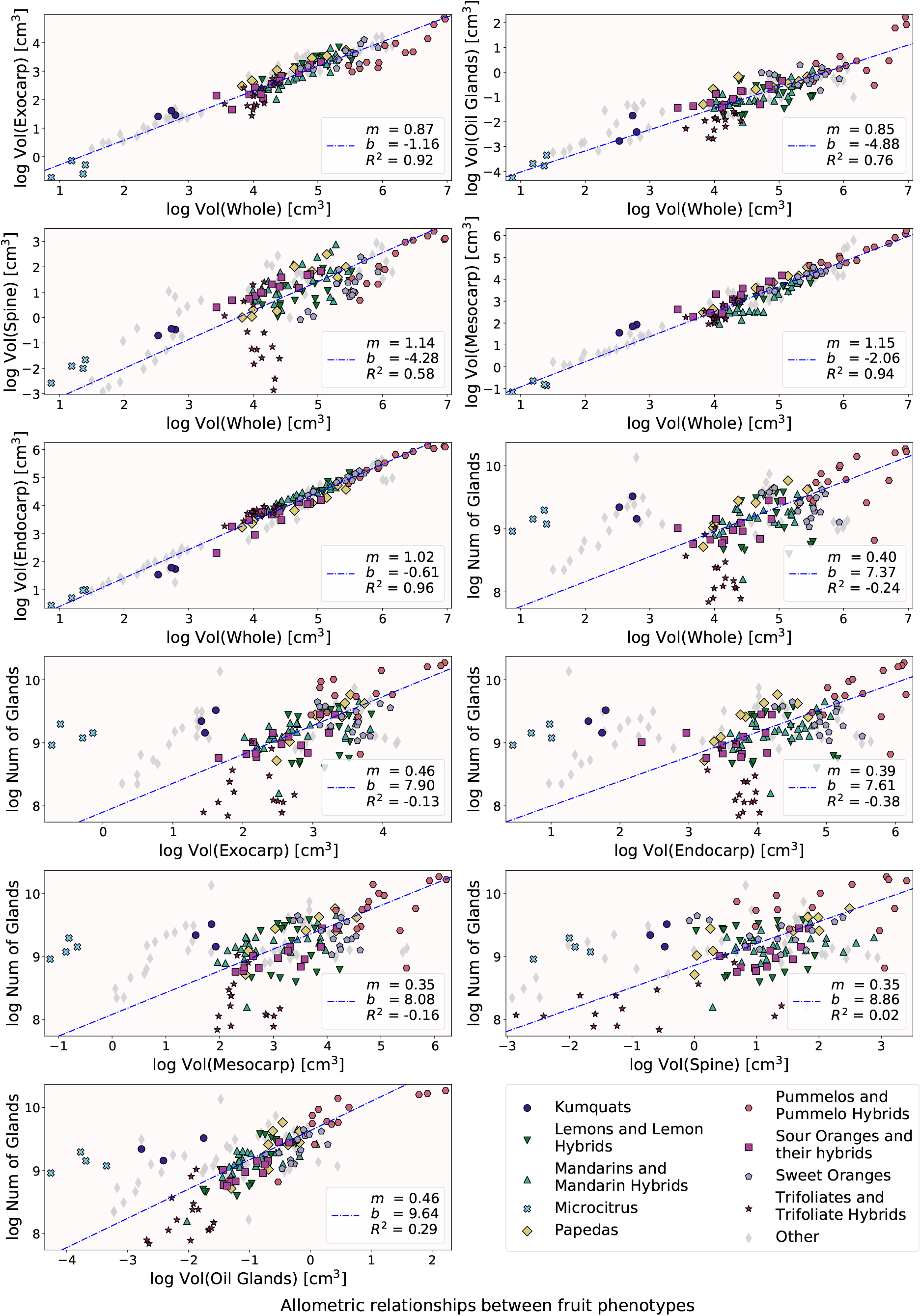
Various allometry plots. We observe a strong allometric relationship between different tissue volumes compared to the total fruit volume across all fruits. The best fit line is depicted by a dashed line in blue. For each plot, the slope, intercept, and correlation coefficient are recorded as *m, b*, and *R*^2^ respectively. The linear relationship in the log-log plots suggests that fruit tissues may grow following a power law. 11

**FIGURE 3.**
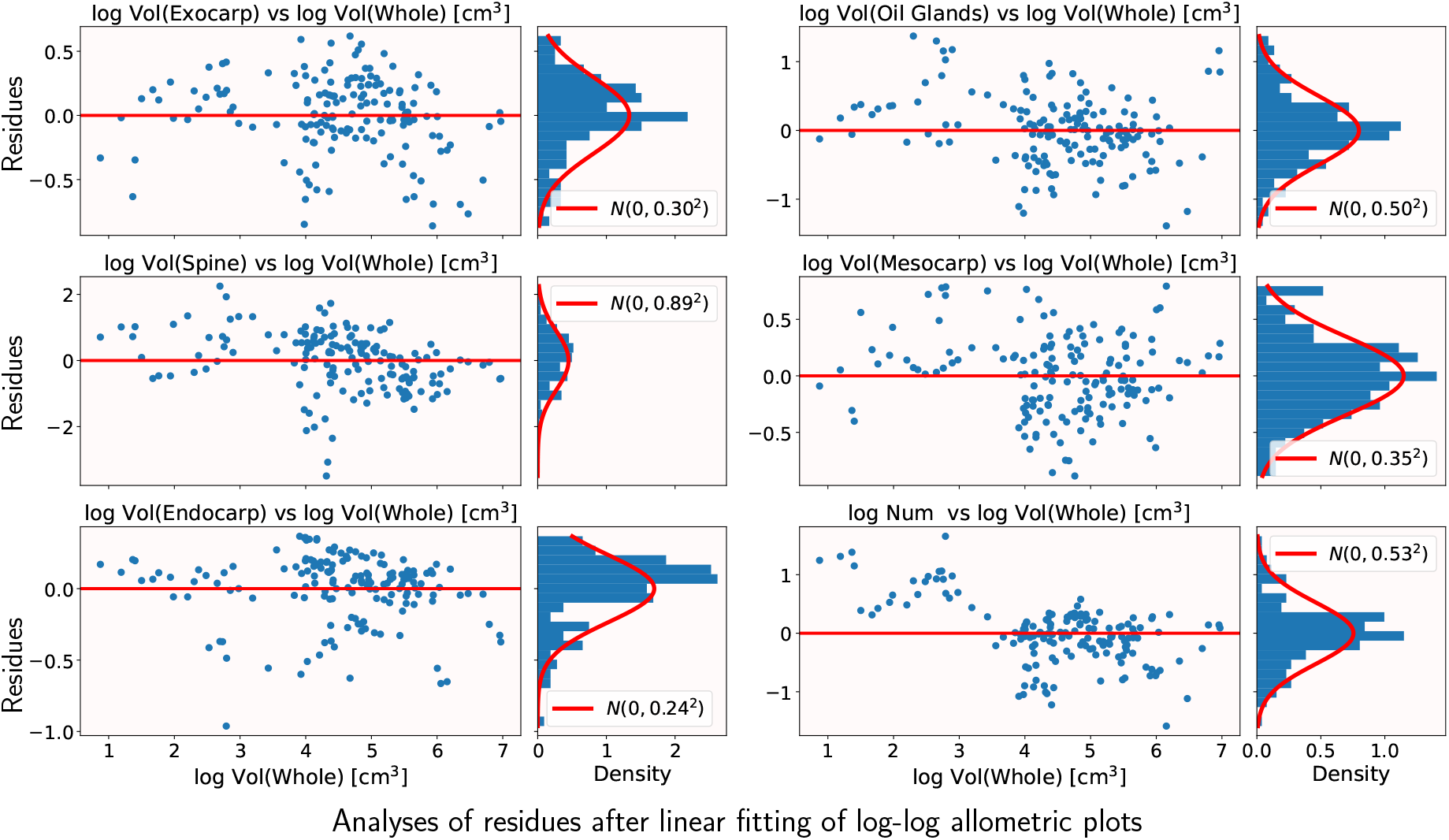
Analysis of the distribution of residues. Analysis of the distribution of residues. The left side of each column are the residues of the fitted linear regression from Figure 2. The right side of each column shows the distribution of these residues. For some of these measurement pairs of traits, the residues follow a normal distribution, suggesting that the linear fit in the log-log plots is adequate.

### 2.3 Oil gland distribution

For each fruit, a point cloud, a collection of (*x, y, z*) coordinates in the space, was defined by the centers of all its individual oil glands. The 25 nearest neighbors, based on Euclidean distance, were computed for each point, so that distances are not affected by the fruit skin curvature. The average distance between each gland and its nearest neighbor, its second nearest neighbor, and so on were computed. The oil gland density was determined both in terms of volume and surface area, by dividing the number of glands by the volume of the whole fruit, and by the surface area of the best-fit ellipsoid (discussed later in Section 2.4) respectively. As in the previous section, all these measurements were log-transformed, linear regressions fitted, parameters recorded, and residues compared to a normal distribution (Figures 4(a), S3). A root square model was fitted between the average nearest neighbor distance and the nearest neighbor index to describe how far apart glands spread from each other (Figure 4(b)).

**FIGURE 4.**
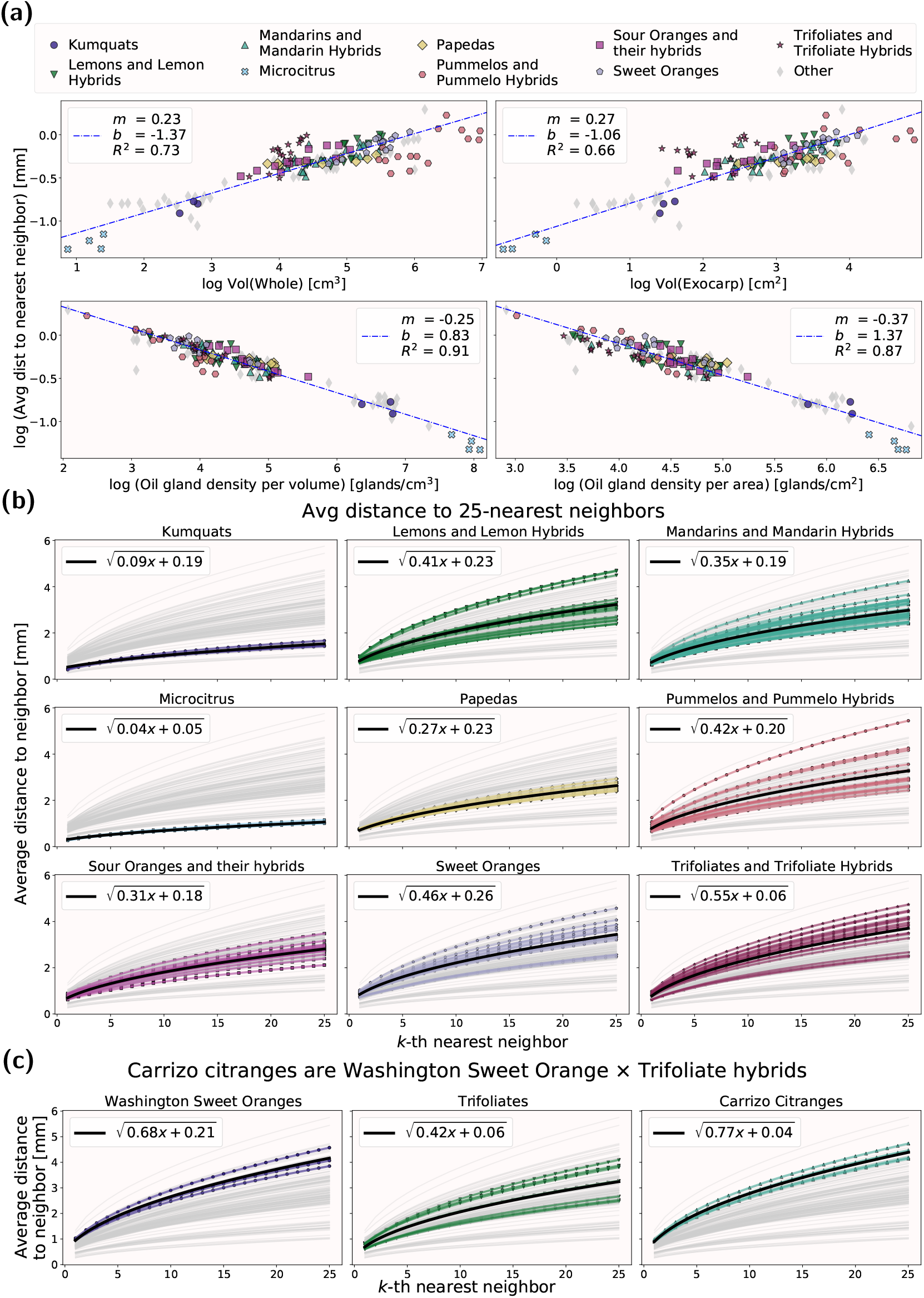
Studying the average distance from each gland to its nearest neighbors. (a) Allometric relationships are observed across all fruits when comparing the average distance between each gland to its closest neighbor with the overall size of the fruit. The overall linear trend is depicted by a dashed blue line. The slope, intercept, and correlation coefficient are denoted by *m, b*, and *R*^2^ respectively. (b) For each oil gland, the average distance to its nearest neighbors follows a square root relationship. The average for each group is plotted as black, thick line. These square root models follow different parameters depending on the citrus group. The fruits that deviate the most from the average usually correspond to hybrids. (c) Carrizo citranges are Washington sweet orange × Trifoliate hybrids. The average distance between oil glands increases at a faster rate for citranges than for their parents, which suggests hybrid vigor.

### 2.4 Modeling the whole fruit as an ellipsoid and computing its sphericity

The surface of most of citrus fruits and their relatives can be approximated by an ellipsoid, a sphere with its three main axes possibly shrunk or stretched.The three axes of symmetry of an ellipsoid delimit three line segments from the center of the ellipsoid to its surface. These are referred to as the *ellipsoid semi-axes*. Notice that a sphere is an ellipsoid with its three semi-axes of the same length. We will consider triaxial ellipsoids, where the length of each semi-axes can be different. An ellipsoid can also be represented as a quadratic equation surface which is both mathematically simple to manipulate (Harris and Stöcker, 1998, Ch. 8.12), and versatile enough to represent both the shapes of nearly-spherical Valencia oranges and elongated finger limes given the right semi-axes lengths.

Each fruit is defined by a point cloud made by the centers of all its individual oil glands. The parameters of the best-fit ellipsoid for this point cloud are computed following the algorithm by Li and Griffiths (2004), from which the semi-axes lengths, rotations, and center are determined (Panou et al., 2020). The fruit point cloud is then rotated and translated such that the best-fit ellipsoid is centered at the origin and its semi-major axes coincide with the proximal-distal axis of the fruit. Finally, the centers of the oil glands are projected to this ellipsoid via geocentric projection, where a ray from the center of the ellipsoid to the gland is drawn and its intersection with the ellipsoid is considered (Figure 5(a)–(f)).

**FIGURE 5.**
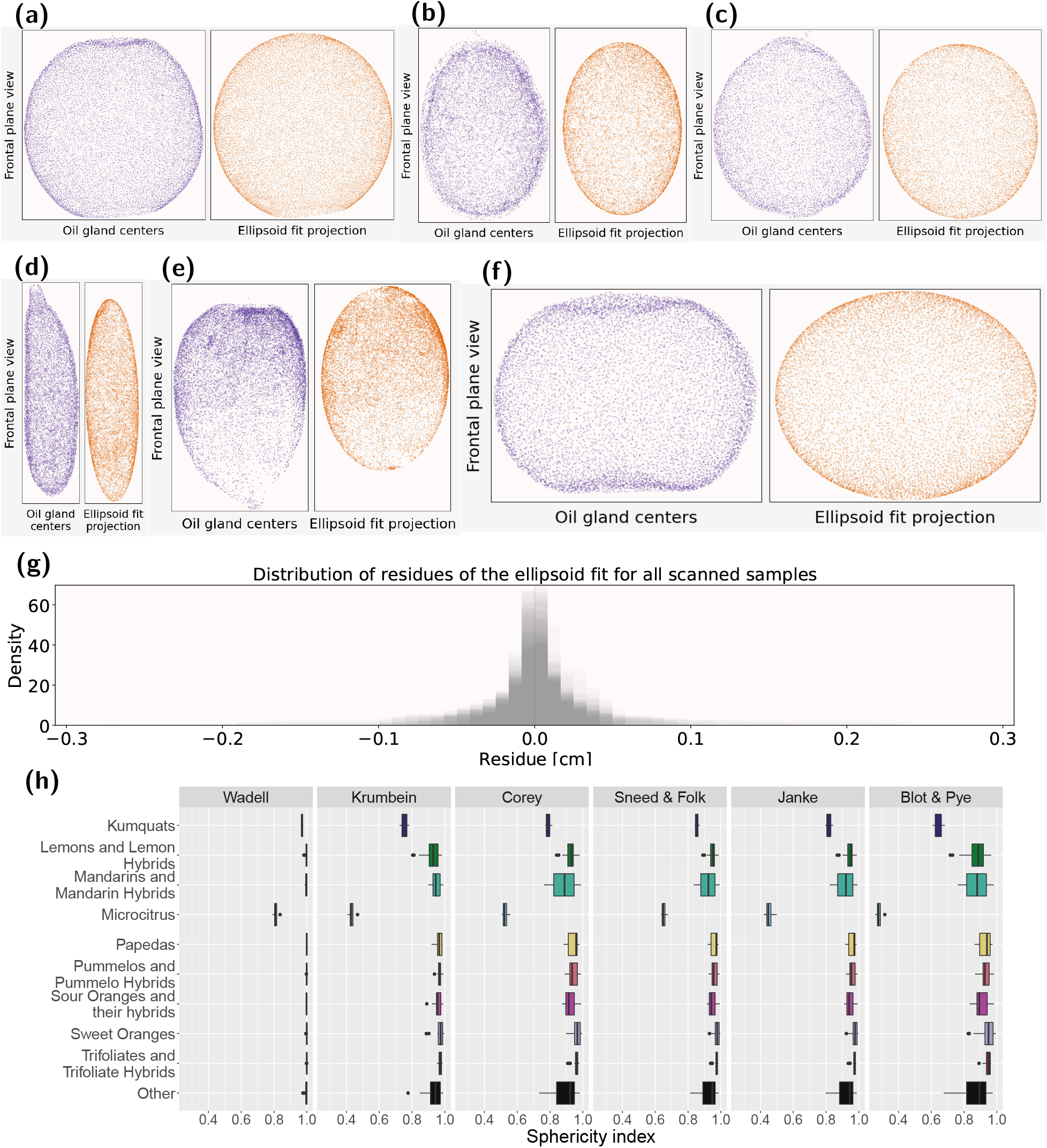
Modeling citrus as ellipsoids. The surface of different citrus was approximated with tri-axial ellipsoids. The glands were later centered at the origin, and the ellipsoid aligned with the proximal-distal, medial-lateral, and adaxial-abaxial axes. Next, the oil glands were projected to this best-fit ellipsoid simply by projecting a ray from the origin —also known as the geocentric projection. Examples of a (a) Cutter nucellar Valencia orange, (b) Nagami kumquat, (c) Willowleaf sour orange, (d) Australian finger lime, (e) South Coast Field Station citron, and a (f) Cleopatra mandarin. Figures (a)–(f) are not scaled. (g) Distribution of the residues of the centers of the oil glands to the best-fit ellipsoid. The distributions for all the fruit scans are overlaid. (h) Various sphericity indices are computed and compared across different citrus groups. The indices are named according to their original reference in Table 2.

This ellipsoid model summarizes important information of the overall shape of the fruit. As an example, we measure how sphere-like different citrus are. There is no unique way to measure sphericity, however, most of the commonly used formulas are based on the semi-axes lengths of the object (Blott and Pye, 2008; Clayton et al., 2009). We measured the sphericity of the resulting fruit-based ellipsoids using 6 different sphericity indices, all of them taking values between 0 (planes and lines) and 1 (perfect spheres) (Figure 5(g); Table 2).

**TABLE 2.**
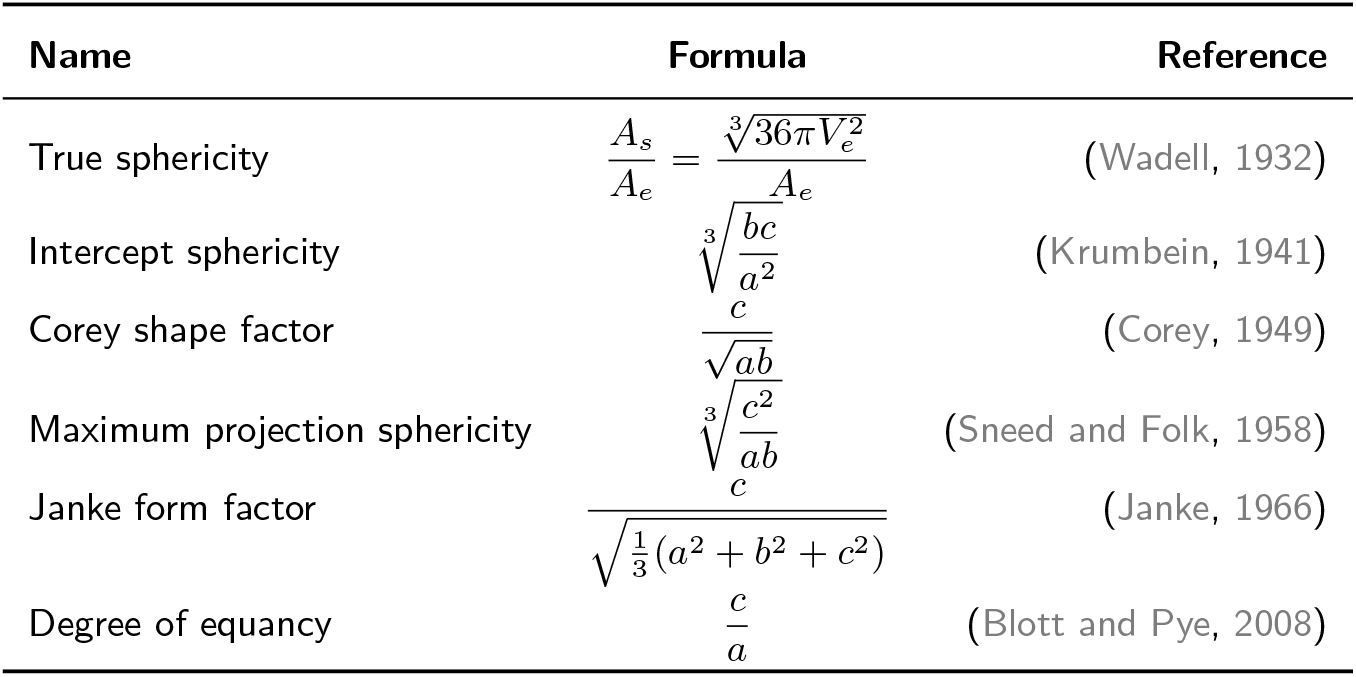
Common sphericity indices based off best-fit ellipsoid. The indices values are bounded between 0 (line or plane) and 1 (perfect sphere). The surface area, volume, and the largest, intermediate, and smallest semi-axes lengths of the ellipsoid are denoted by *A*_*e*_, *V*_*e*_, *a, b, c* respectively. Also, *A*_*s*_ denotes the surface area of a sphere of volume *V*_*e*_.

### 2.5 Revisiting the distribution of the oil glands

The projected gland center locations on the ellipsoid were described in terms of longitude and latitude coordinates with respect to the ellipsoid (Diaz–Toca et al., 2020). We tested whether the gland point cloud follows a uniform distribution, where every unit area of the skin has the same probability of containing oil glands; or if the underlying distribution is rotationally symmetric, where the oil glands pattern is symmetrical around a fixed direction. Uniformity was tested with Projected Anderson-Darling (PAD) test (García-Portugués et al., 2020) with the R package sphunif (García-Portugués and Verdebout, 2021). The rotational symmetry was tested with a scatter-location hybrid test with an unspecified direction of symmetry (García-Portugués et al., 2020) with the R package rotasym (García-Portugués et al., 2021). Additionally, we visually examined the distribution of oil glands for most fruits and compared them to simulated uniform distributions by projecting them to 2D via Lambert azimuthal equal-area projections (Mardia and Jupp, 1999, Ch. 9.1) from the North and South poles. Intuitively, these two projections flatten the sphere on a plane by pushing it from the North pole and South pole while minimizing the distortion seen on the north and south hemisphere respectively (Figure S5).

## 3 Results

### 3.1 Allometric relationships

The estimated volume of each tissue type and fruit follows the expected average fruit size of each genetic group, with the smallest fruit in the bottom left corners (microcitrus, kumquats) and large fruit in the top right corners (pummelos). Strong linear trends are observed when comparing most of the volume-related features of all the fruits, indicated by high *R*^2^ correlation coefficient values, usually above 0.75 except for the central column tissue (Figures 2, S1). Due to their thin size and scanning quality, these columns were difficult to identify and isolate, especially in some trifoliates, which might explain lower *R*^2^ values. The residues of the fitted linear regression tend to follow a normal distribution for the majority of measurement pairs, suggesting that the linear fit is adequate (Figures 3, S2). This linearity indicates that the tissues across all citrus fruits grow relative to each other following a power rule. For example, looking at the slope *m* values, both the exocarp and the oil glands grow in volume at the same relative rate with respect to the volume of the whole fruit (*m* = 0.85). On the other hand, the total number of oil glands appears to be decoupled from all the measured size-related traits, as shown by much lower *R*^2^ values (Figure 2). In this case, a power law may not be an adequate model to describe the oil gland number with respect to tissue volume.

### 3.2 Oil gland distribution

There is a strong positive linear relationship between the volume of the fruit, and the average distance between an oil gland and its nearest neighbor, with *R*^2^ correlation coefficients above 0.65. There is a stronger negative linear relationship when considering the overall oil gland density, reflected by *R*^2^ coefficients above 0.85 (Figure 4(a)). The residues follow normal distributions, indicating that the linear model is adequate (Figure S3). These allometric relationships suggest that for all citrus and relative fruits, the average distance between nearest oil glands follows a power law with respect to fruit volume and gland density. When considering fruit size, as expected, the samples distribute in a similar pattern as with most of the previous allometry plots. However, an inverse pattern is observed when considering oil gland density. In this case, the smallest fruits tend to report the highest number of oil glands per unit volume or unit area. Other than microcitrus and kumquats, the rest of highlighted citrus groups form a tighter cluster. The average distance between an oil gland and its *k*-th nearest neighbor is modeled as

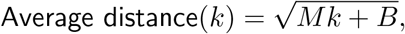

where *M* is the rate of distance growth and *B* the line intercept (Figure 4(b)). As expected from their higher oil gland density, the average distance to the gland’s nearest neighbors increases the slowest for microcitrus, followed distinctly by kumquats. On the other hand, sweet oranges and trifoliates report the largest average distances between neighboring oil glands.

In general, all the samples of every accession follow the same growth model. Outliers in growth models are typically associated with hybrid accessions. Increased growth rates are found when the second parent is a large fruited accession. For example, consider the Carrizo citrange, a trifoliate x Washington sweet orange hybrid where sweet orange fruits are much larger than trifoliate fruits. We observed that the oil glands in the citrange grow on average farther apart from each other than in any of the parents (Figure 4(c)), which suggests that hybrid vigor might be at play. Similarly, hybrids derived from crosses with small-fruited accessions have reduced growth rates.

### 3.3 Ellipsoid modeling and sphericity of fruits

The best-fit ellipsoid successfully captures the overall shape of the citrus and relatives, with a negligible portion of gland centers differing by more than 0.2cm from their ellipsoidal approximation (Figure 5(g)). This ellipsoidal model is flexible enough to capture both spherical and elongated fruit shapes, from sweet oranges to Australian finger limes (Figures 5(a)–(f)).

Most of the fruits report highly spherical indices, more than 0.9 for every sphericity index. Unsurprisingly, the elongated Australian finger limes are the least spherical and their shape is very distinct from the rest of the samples. The kumquats are less elongated than the finger limes, but they also remain highly distinguishable for most of the sphericity indices. Mandarins and their hybrids also tend to be slightly less spherical than the remaining groups of interest (Figure 5(h)).

### 3.4 Oil glands revisited

Although the tools from directional statistics assume that the data points lie on a sphere rather than an ellipsoid, most of scanned fruits were very sphere-shaped according to a variety of sphericity indices (Figure 5(h)). Thus shape information is not significantly altered when translating longitude and latitude coordinates from the best-fit ellipsoid to a sphere.

The uniform oil gland distribution hypothesis was strongly rejected for all scans, with all p-values below 0.015 for the PAD test, and below 2.5 × 10^−7^ for 95% of all the point clouds (Figure S4(a)). The scatter-location hybrid test strongly rejected the rotationally symmetric oil gland distribution hypothesis for most of the point clouds as well. More than 90% of all the scans reported p-values smaller 0.02. The 10 samples for which the rotationally symmetric hypothesis was not rejected were not concentrated in any citrus groups (Table S3; Figure S4(b)). Upon closer visual examination, differences arise between the uniform distribution on a sphere and oil gland distributions. The oil gland distributions tend to have defined clusters and empty spots, which are not seen in typical uniform distributions. The northern and southern hemispheres are noticeably different from each other for the oil gland distributions, while these look roughly the same in uniform distributions (Figure S5).

## 4 Discussion

Measuring and understanding the shape is fundamental to extracting valuable information from data, and push further our insights. A vast number of biological-inspired shapes are intrinsically 3 dimensional, like citrus fruit, and capturing their shape as 3D voxel-based provides a faithful shape representation that allows accurate measurement of tissue volumes and modeling of gland distributions in space. Better fruit modeling is key to provide more accurate descriptions of fruit shape and oil gland content, as both are important traits for citrus scion improvement (Barry et al., 2020). Citrus shape impacts oil gland abundance and distribution, as the shape of the rind, the exocarp, and other tissues, along the distribution of the oil glands affects the physics of citrus essential oil extraction and aroma dispersion (Smith et al., 2018).

When observing overall fruit tissue size trends, these correspond to known citrus genealogy. For example, when comparing the size of exocarp against size of endocarp, most of the sour and sweet oranges tend to lie between mandarins and pummelos, with sour oranges lying closer to mandarins, while the sweet oranges are closer to pummelos (Figure 2). A similar arrangement of citrus groups is observed when comparing the average distance between neighboring oil glands to either fruit volume or gland density. Moreover, it is observed that oil glands distance themselves from each other following a square root rate in general. The exact degree to which they push each other apart depends on the oil gland density, which in turn is partly affected by the citrus genealogy. For example, the average distances between oil glands in Carrizo citranges increase at higher rates than in either the Washington sweet oranges or the trifoliates, the citrange parents (Figure 4). The square root suggests that the mechanics of oil gland displacement across the fruit could be partly governed by based on Brownian motion and normal diffusion interactions (Vlahos et al., 2008). However, the hypothesis of oil gland locations following either a uniform or symmetric distribution on the fruit surface is strongly rejected for all scans by the PAD and location-scatter hybrid tests respectively. This discrepancy between our gland distribution observations could be explained by the fact that uniform distributions and diffusion processes assume that the data consists of point particles with no volume that can stand arbitrarily close to each other. For oil glands this is obviously not the case, as they have volume and there are physical limitations on the proximity between glands, which requires a more complex diffusion modeling. Higher resolution scans might be able to capture better individual oil gland shape, rather than just its center. Individual oil glands then could be approximated by individual minimum volume enclosing ellipsoids (Todd and Yildirim, 2007), which could then pose more advanced distribution and diffusion models. At the same time, more advanced tools from directional statistics, like spherical kernel density estimators (Di Marzio et al., 2019; Vuollo and Holmström, 2018) might be able to characterize the oil gland distributions in a non-parametric way, while allowing a more numerical comparison of their differences among distinct citrus groups.

All the studied citrus and related accessions exhibit allometric behavior in general across both their tissue volumes, and average distances between neighboring glands. This relative growth relationships suggest that tissue sizes are deeply linked, as the size of oil gland tissue in general may not be able to change without changing volume of both the endocarp and mesocarp. Moreover, there might be biophysical principles at play that govern different tissue development across all citrus fruits in general, just like normal diffusion might govern oil gland distribution. The determination of such biophysical constraints prompt future lines of exciting research.

There is rich shape information in the natural world to be captured, analyzed, and linked to biophysical developmental and evolutionary principles. Sound mathematical models are key to uncover these biophysical interactions at work. For example, even with a limited number of points, overall fruit shapes can be approximated with various quadratic surfaces like ellipsoids. Given the appropriate parameters, an ellipsoid can represent both nearly spherical navel oranges, and elongated finger limes. This quadratic surface approximation is mathematically versatile and computationally simple, and can be applied to other round-shaped biologically-motivated data. Moreover, ellipsoid coordinates can be translated naturally to longitudes and latitudes on a sphere, which opens the door to a wide array of mathematical tools from directional statistics. Some of those tools, like density estimations, hypothesis testing, and distribution fitting, allow us to quantify shape in a mathematically rigorous and comprehensive way. The quality of input data for our models is equally important. Through X-ray CT scanning technology we have a novel way to observe, quantify, and analyze all the shape of citrus and their tissues in a comprehensive, automated, non-invasive, and non-destructive manner. With the right voltage and current, the 3D X-ray CT reconstructions can discern small, individual tissues, like oil glands, which enables us to analyze tissue shape and distribution at very granular levels. Capturing and analyzing this nuanced shape information for a wide array of data sets provides a morphology-driven path to further our insight into phenotype-genotype relationships. As stated by D’Arcy Thompson in his seminal biomathematical treatise *On Growth and Form*, “An organism is so complex a thing, and growth so complex a phenomenon, that for growth to be so uniform and constant in all the parts as to keep the whole shape unchanged would indeed be an unlikely and an unusual circumstance. Rates vary, proportions change, and the whole configuration alters accordingly” (1942).

## Acknowledgements

Daniel Chitwood is supported by the USDA National Institute of Food and Agriculture, and by Michigan State University AgBioResearch. The work of Elizabeth Munch is supported in part by the National Science Foundation through grants CCF-1907591, CCF-2106578, and CCF-2142713.

## Author contributions

EA, DS, EM, and DC conceived the experiment. DS selected the accessions to be scanned and collected the plant material, ensuring that citrus types were broadly represented. MQ and DC collected the digital data. EA and TO developed the necessary scripts to process the scans and extracted their shape descriptors. EA analyzed the data and wrote the manuscript. All authors contributed, reviewed, and revised the manuscript.

## Software and data availability

The processed and cleaned citrus X-ray CT 3D reconstructions can be found in the Dryad repository https://doi.org/10.5061/dryad.34tmpg4n6, along with their separated tissues and associated point clouds and ellipsoidal approximations.

All our code is available at the https://github.com/amezqui3/vitaminC_morphology_repository. This includes the image processing pipeline to clean the raw scans and segment the fruit tissues, the computation of tissue volume, the best-fit ellipsoid, and the hypothesis testing of uniform and symmetric distributions on a unit sphere. A collection of Jupyter notebook tutorials is also provided to ease the usage and understanding of the different components of the data processing and data analyzing pipelines. All the image-related scripts are available in python, while the statistical analyses are in R.

## Supplementary data

**FIGURE S1.**
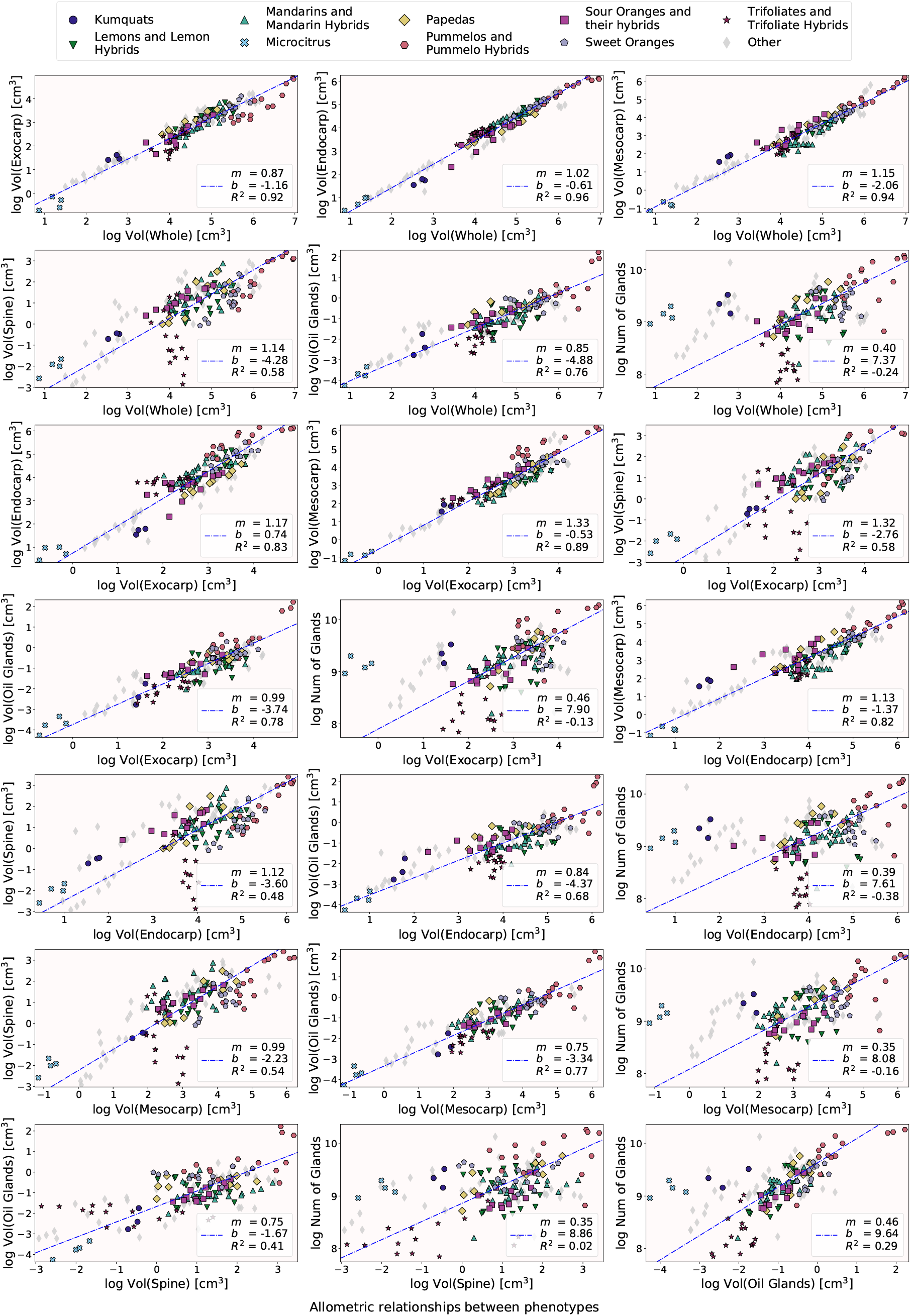
Allometry plots for all possible pairs of measured phenotypes. We observe a strong allometric relationship between different tissue volumes. However, this relationship is missing when comparing the total number of oil glands to all tissue volumes, suggesting that the number of glands is decoupled from these volume traits. The best fit line is depicted by a dashed line in blue. For each plot, the slope, intercept, and correlation coefficient are recorded as *m, b*, and *R*^2^ respectively. The linear relationship in the log-log plots suggests that fruit tissue volumes may grow following a power law.

**FIGURE S2.**
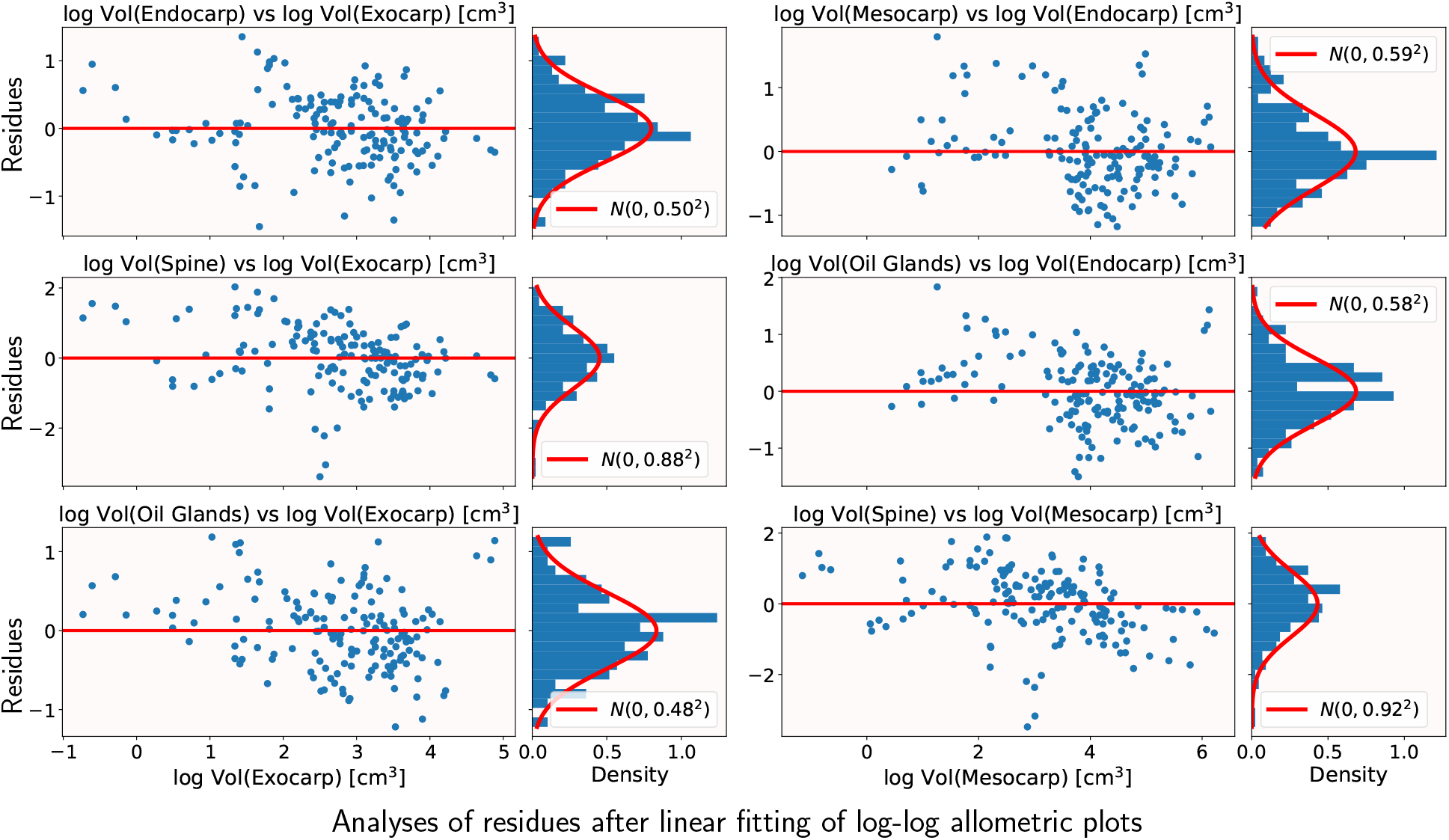
Analysis of the distribution of residues as in Figure 3. The left side of each column depicts residues of the linear fit as in Figure S1. The right side of each column shows the distribution of these residues. For some of these measurement pairs of traits, the residues follow a normal distribution, suggesting that the linear fit in the log-log plots is adequate.

**FIGURE S3.**
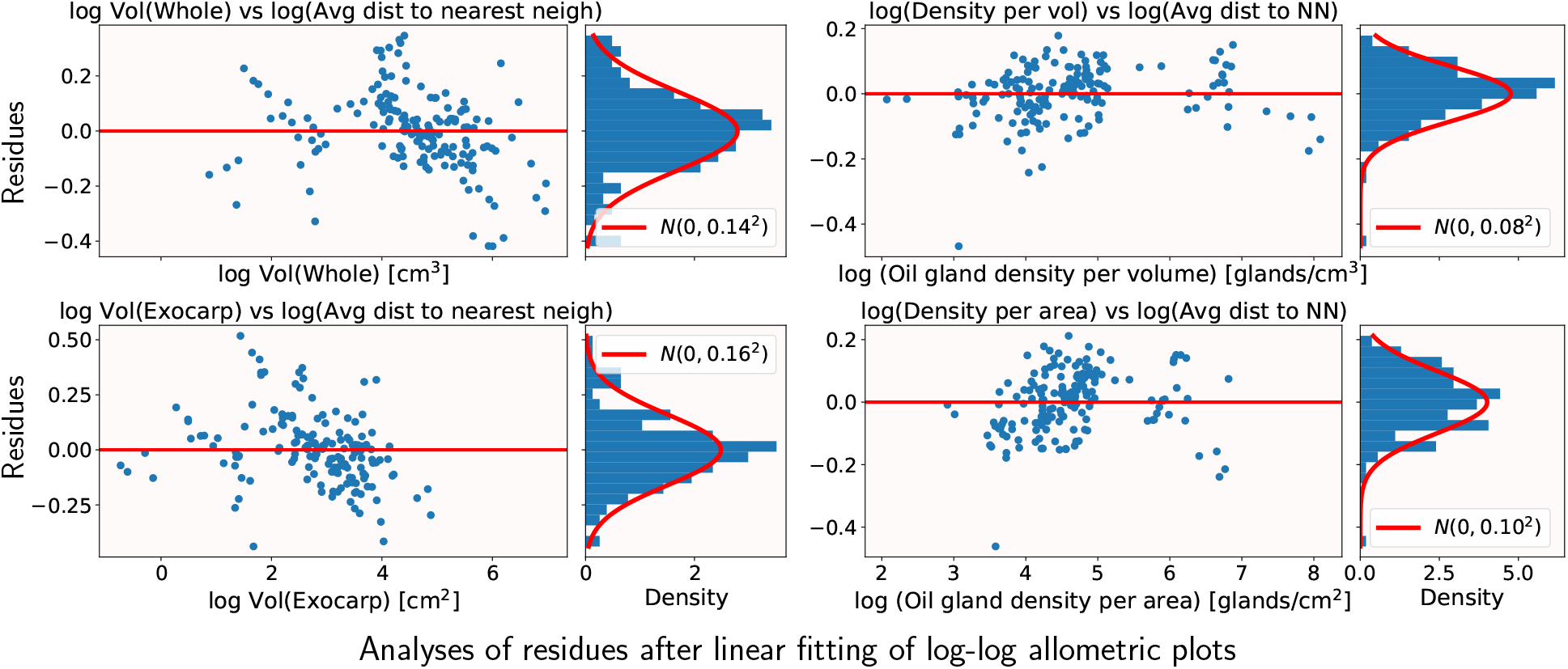
Analysis of the distribution of residues. The left side of each column depicts residues of the linear fit as in Figure 4(a). The right side of each column shows the distribution of these residues. For some of these measurement pairs of traits, the residues follow a normal distribution, suggesting that the linear fit in the log-log plots is adequate.

**FIGURE S4.**
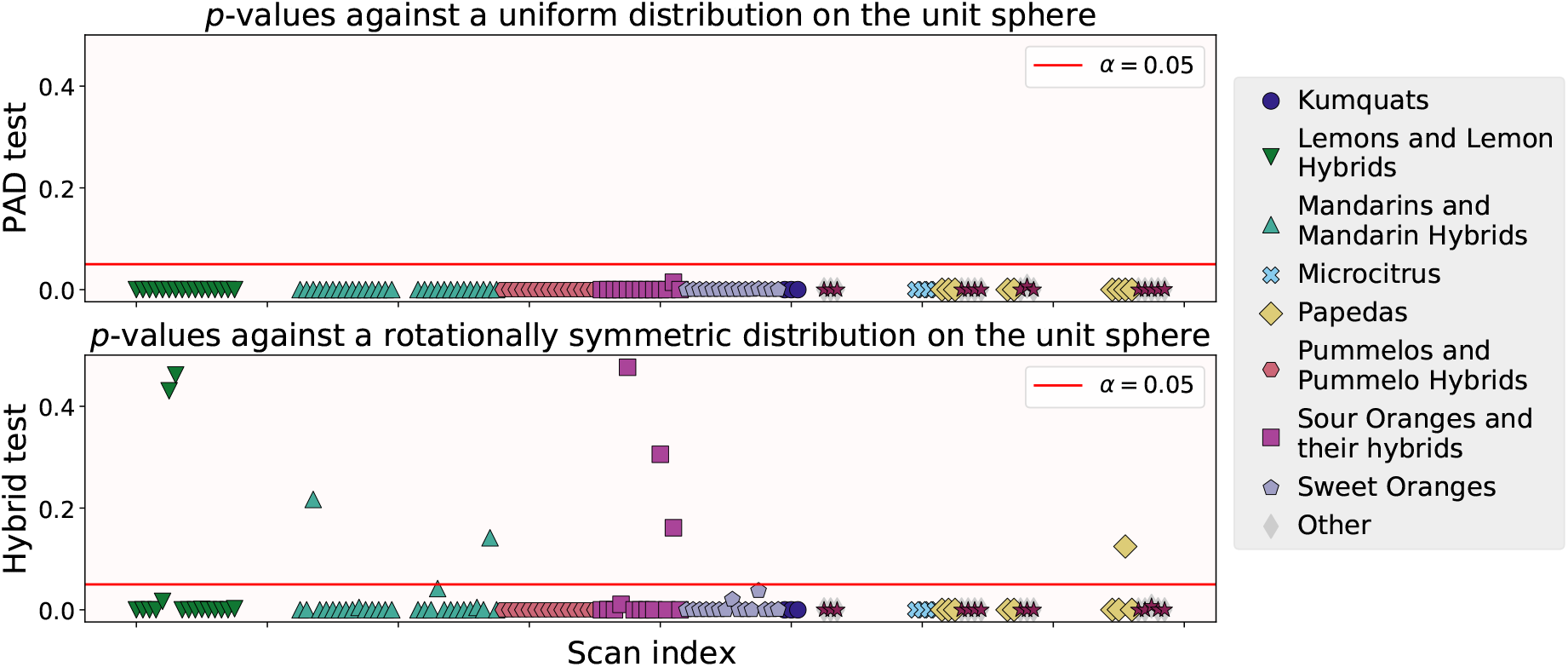
Testing whether the oil glands are distributed uniformly or rotationally symmetric on the surface of the fruits. (a) *p*-values after testing if the underlying oil gland distribution is not uniform according to projected Anderson-Darling (PAD) test. (b) *p*-values after testing if the underlying oil gland distribution is not rotationally symmetric according to the scatter-location hybrid test. Red line at *α* = 0.05 in all plots.

**FIGURE S5.**
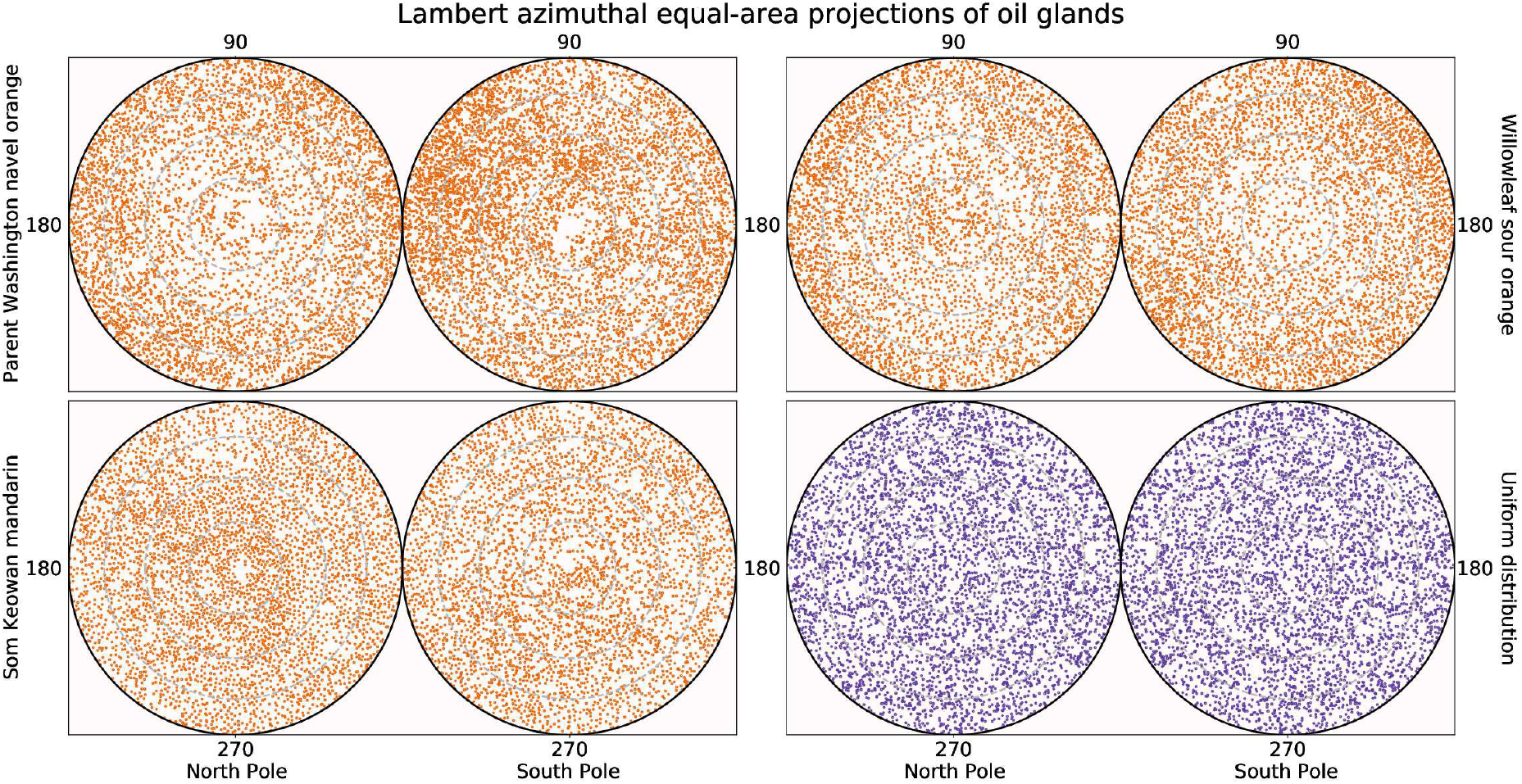
Distribution of oil glands is not uniform across the citrus exocarp. After modeling the fruits as ellipsoids and projecting their oil glands onto the ellipsoidal surface, longitude and latitude coordinates are computed as in Figure 5. These coordinates can be better visualized using two Lambert azimuthal equal-area projections, from the north and south poles which represent the northern and southern hemispheres respectively with minimal distortion. A battery of statistical tests strongly rejects the hypothesis of these glands being uniformly distributed over the ellipsoid surface. From the top-left corner, examples in orange color are shown of oil gland distribution of a parent Washington navel orange, a Willowleaf sour orange, and a Som Keowan mandarin. For comparison, a similar number of points following a simulated uniform distribution is shown in purple on the bottom-right corner.

