## Supplementary figures and images for "The shape of aroma: measuring and modeling citrus oil gland distribution"

### Supplemental Figure S1

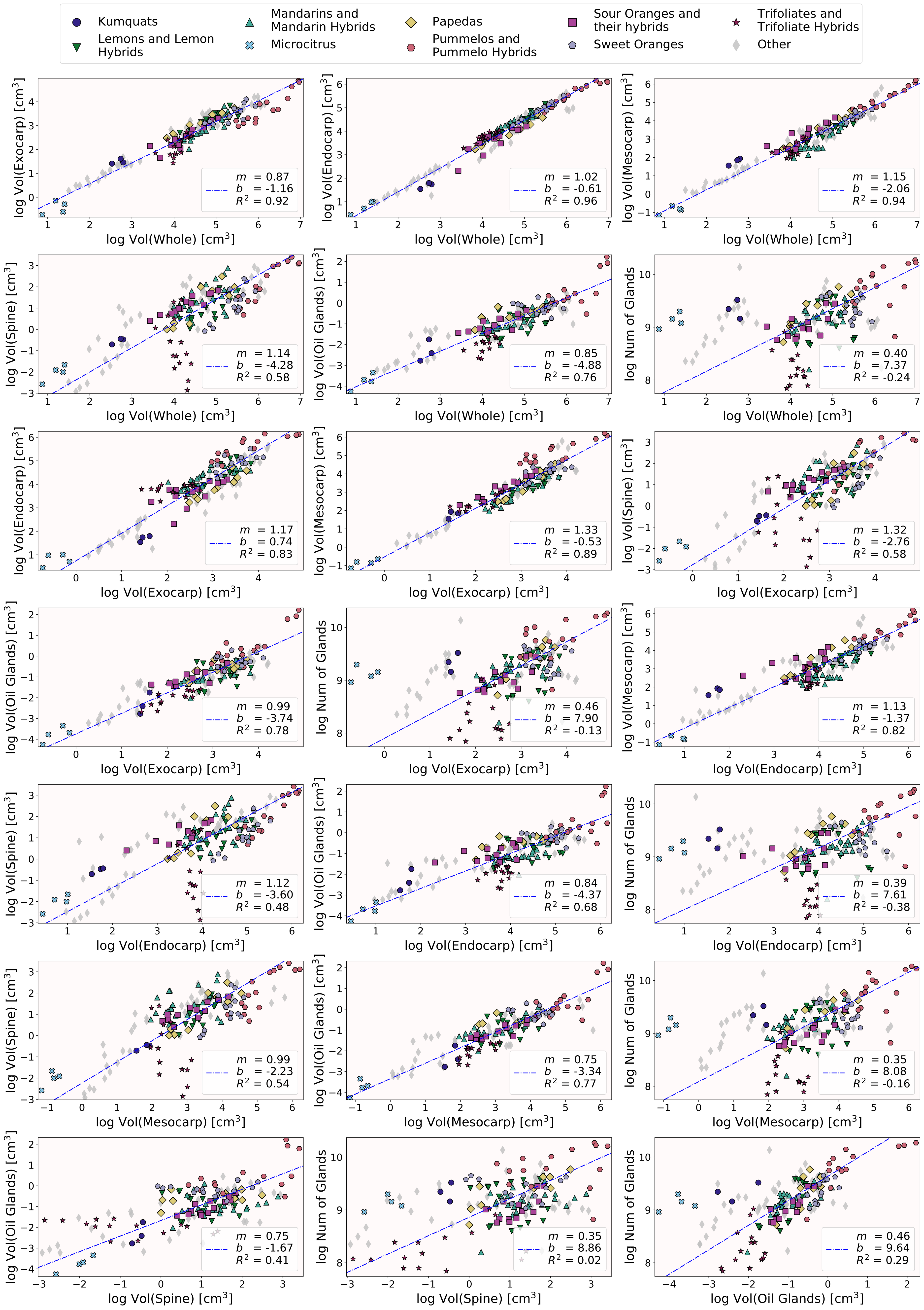


Allometric relationships between phenotypes

### Supplemental Figure S2

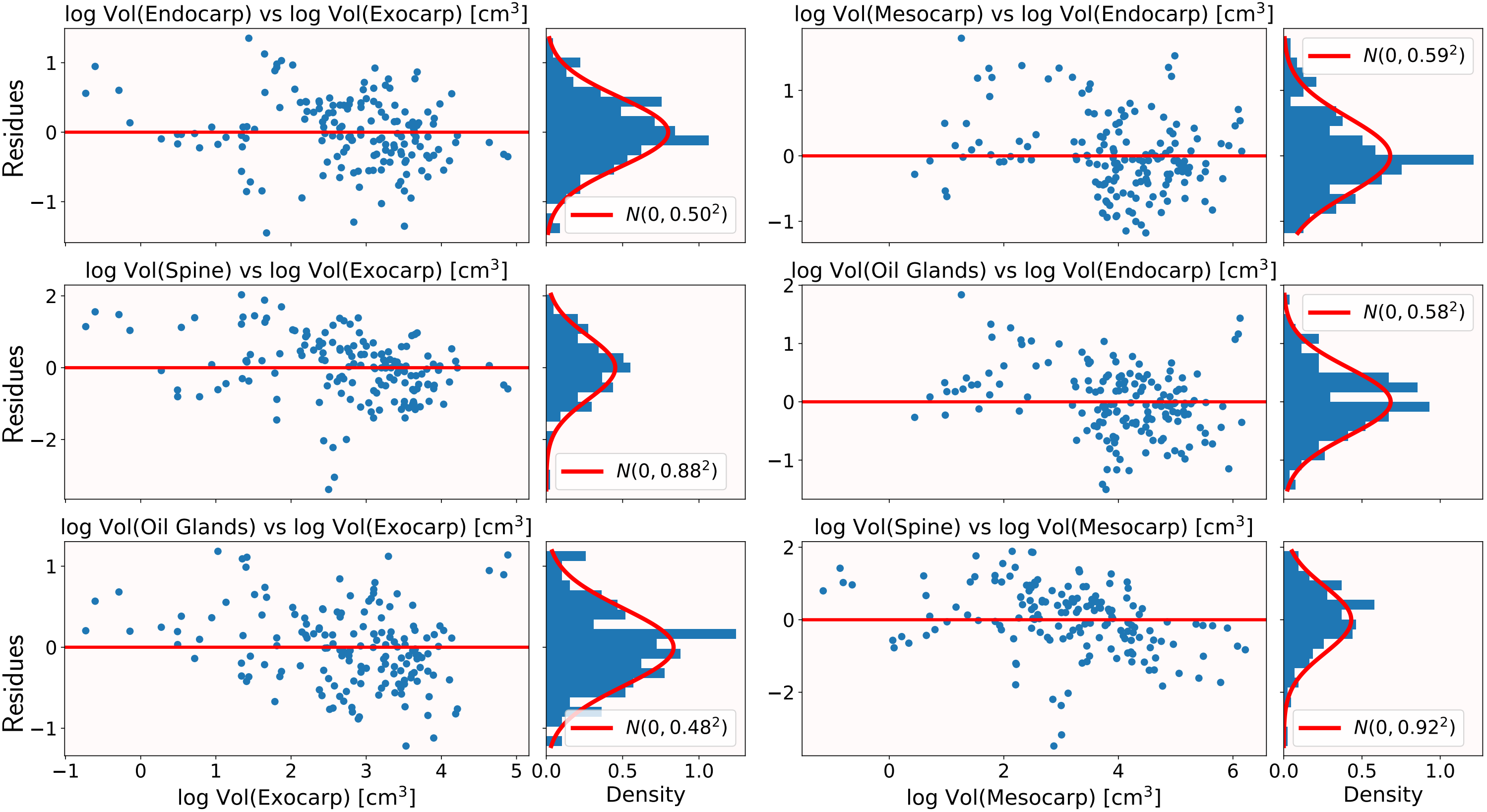


Analyses of residues after linear fitting of log-log allometric plots

### Supplemental Figure S3

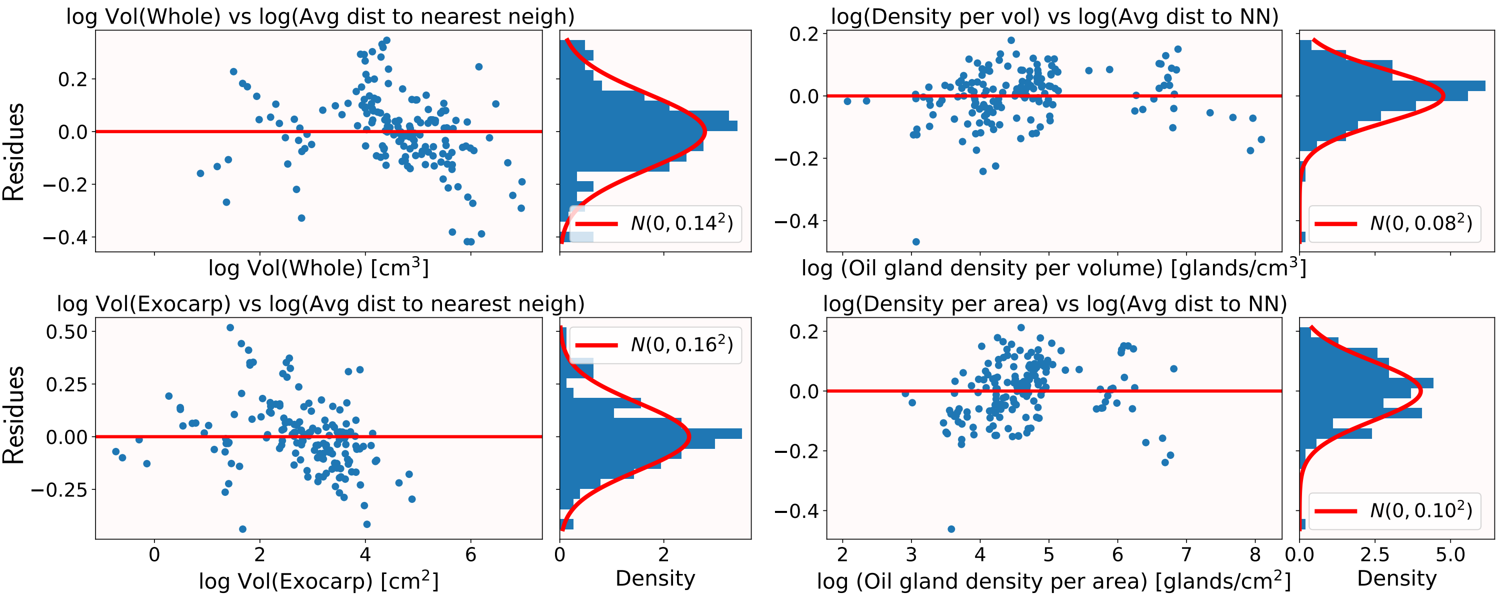


Analyses of residues after linear fitting of log-log allometric plots

### Supplemental Figure S4

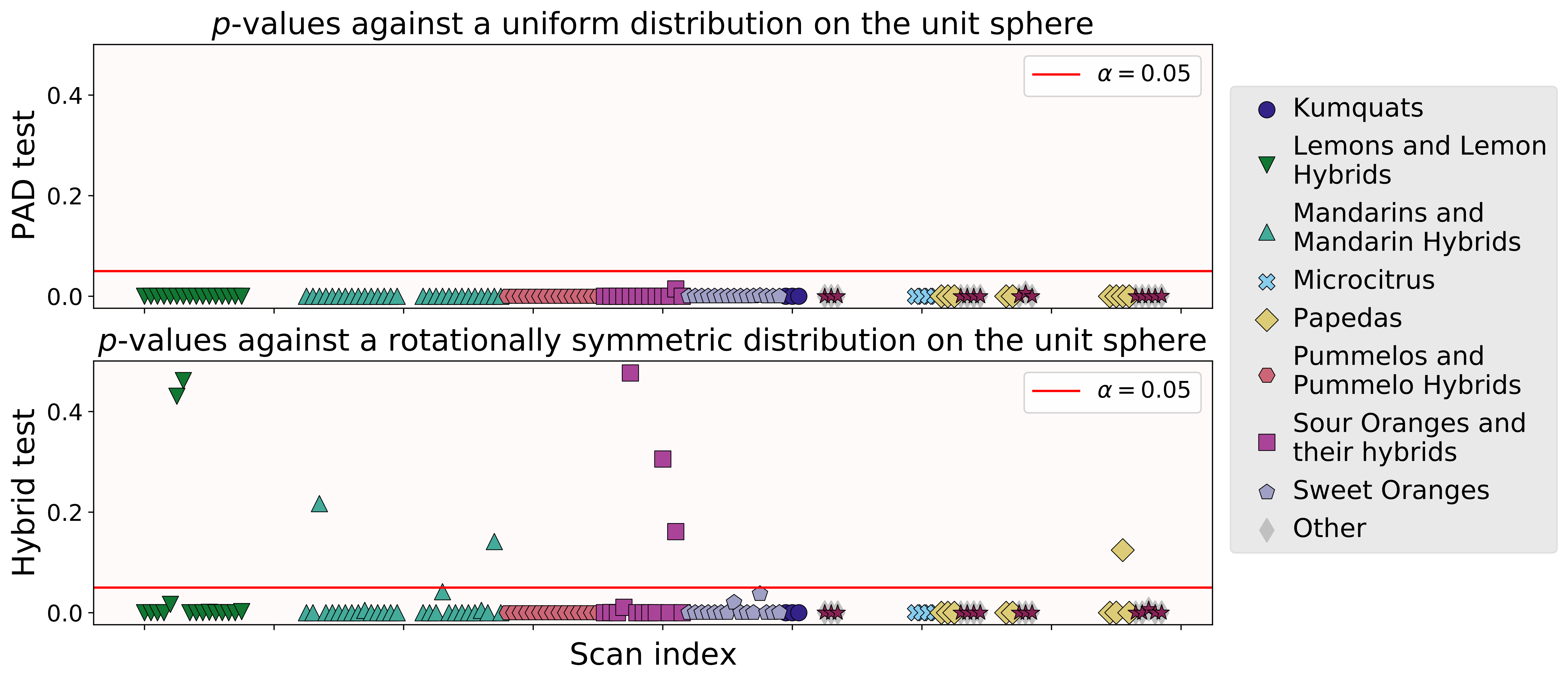

### Supplemental Figure S5

#### Lambert azimuthal equal-area projections of oil glands

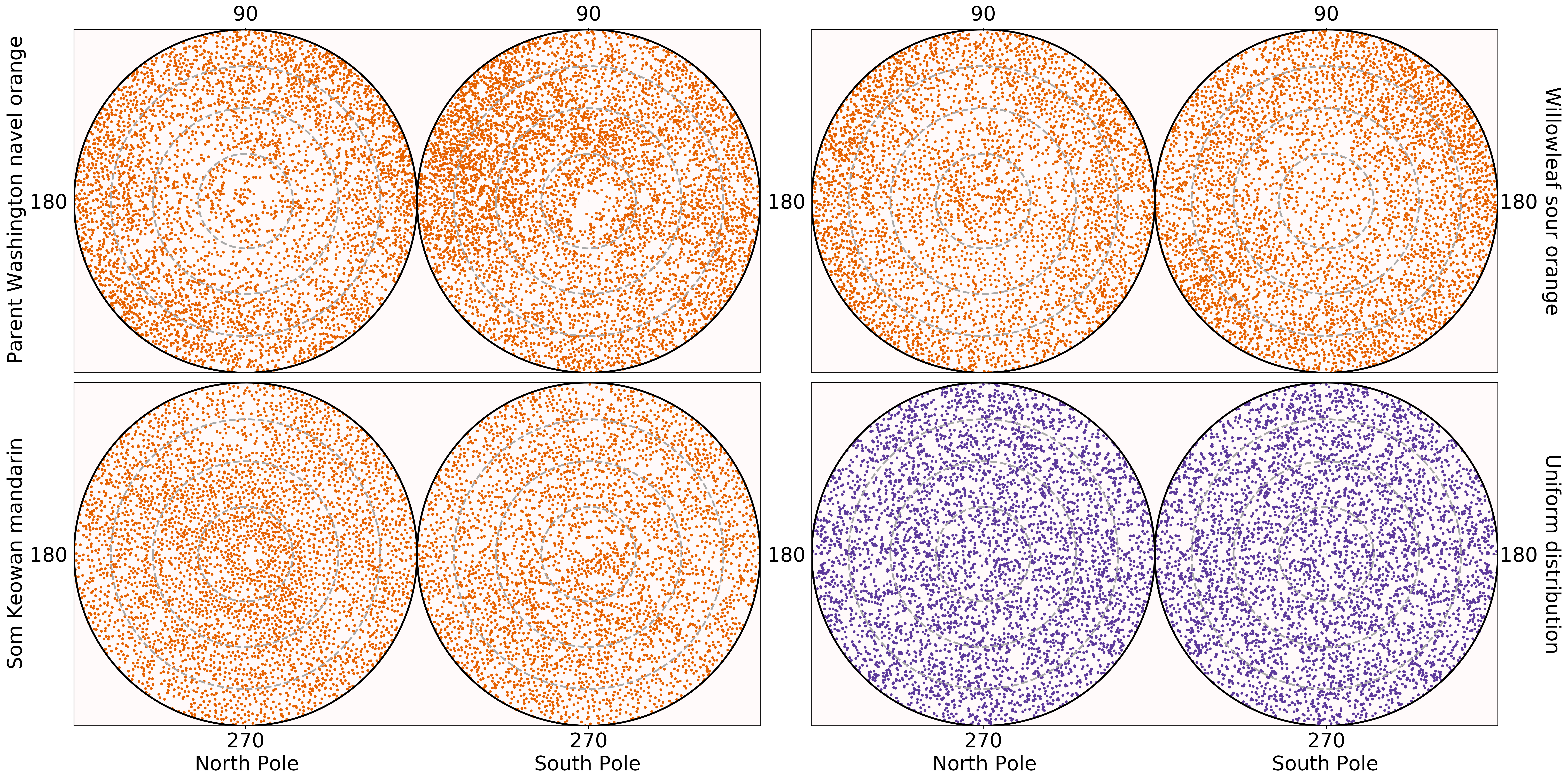
